# Panera: A novel framework for surmounting uncertainty in microbial community modelling using Pan-genera metabolic models

**DOI:** 10.1101/2023.10.11.561816

**Authors:** Indumathi Palanikumar, Himanshu Sinha, Karthik Raman

## Abstract

Over the last decade, microbiome research has witnessed exponential growth, largely driven by the widespread availability of metagenomic data. Despite this influx of data, 16S ‘targeted amplicon’ sequencing, which offers relatively lower resolution, still dominates the landscape over whole-genome shotgun sequencing. Existing algorithms for constructing metabolic models of microbial communities primarily rely on whole-genome sequences and do not fully harness the potential of 16S datasets.

In this study, we report ‘*Panera’*, a novel framework designed to model microbial communities under uncertainty and yet perform inferences by building pan-genus metabolic models. We tested the performance of the models from our approach by analysing their ability to capture the functionality of the entire genus and individual species within a genus. We further exercise the model to explore the comprehensive metabolic abilities of a genus, shedding light on metabolic commonalities between microbial groups. Furthermore, we showcase its application in characterising microbial community models using 16S data. Our hybrid community models, which combine both GSMM and pan-genus metabolic models, exhibit a 10% reduction in prediction error, with error rates diminishing as community size increases.

Overall, the Panera framework represents a potent and effective approach for metabolic modelling, enabling robust predictions of the metabolic phenotypes of microbial communities, even when working with limited 16S data. This advancement has the potential to greatly impact the field of microbiome research, offering new insights into the metabolic dynamics of diverse microbial ecosystems.

## 1 Introduction

The unprecedented growth of metagenomics research in the past decade has shed a spotlight on the significance of microbial communities in various ecosystems encompassing host-associated systems^1–3^, environmental conditions^4^, and even extreme environments like hot springs^5^ and the ocean floor^6^. The composition of microbiomes can be characterised by either amplicon sequencing or shotgun whole genome sequencing (WGS)^7^. While shotgun metagenomics sequences all the genomes present in the sample using random primers, amplicon sequencing targets shorter segments of a genome, most commonly the 16S rRNA gene (16S rDNA), to profile microbiomes. The compositional and functional profiling of a microbiome aids in inferring the existing microbial entities and their activities within an ecosystem^8^. However, it is critical to probe into the intricate interactions within the microbial community as well as with its host and environment.

The reconstruction of genome-scale metabolic models (GSMMs) from genetic, reaction and metabolite information of an organism offers a promising opportunity to comprehend the complex genotype-to-phenotype relations within an organism^9,10^. GSMMs are instrumental in exploring the alteration in physiology and behaviour of the organism under various nutrient conditions and other interventions such as gene or reaction knockouts^11,12^. Prior research analyses have harnessed GSMMs to simulate microbial communities *in silico* and investigate their metabolic dependencies and functionalities^13–16^. However, microbial community modelling has thus far focused only on GSMMs derived from whole genome sequences, simulated as communities based on the precise knowledge of the component individual species in the microbiome. In the context of 16S rRNA, most studies are reported to analyse 250 - 500 bps, only a ⅓ - ⅕ th of the entire gene length (1500 bp)^17^. These short reads constrain the precision of taxonomy assignment, with more than half of the classifications extending only to the genus level^18^. Given the widespread availability of 16S datasets, it is essential to evolve newer frameworks that can leverage this genus-level information to build pan-genus metabolic models (PGMMs) and employ them in the metabolic modelling of microbial communities.

In this regard, it is essential to appreciate the role of pan-genus models in elucidating the unique metabolic capabilities and physiological characteristics of a genus. To date, pan-genus models are either reconstructed using pan genomes or by collecting species-specific GSMM in a single model^13^. Pan-genome-based reconstruction combines the genomes of species within a genus and is used for draft reconstruction using tools such as KBase^19^ and CarveMe^20^, followed by gap-filling and manual curation. For instance, the pan-genome model of *Propionibacterium* assembled from five distinct genomes of the species within this genus has unveiled both the diversity within central carbon metabolism and shared metabolic traits^21^. Similar pan-genome models for *Escherichia*^22^ and yeast^23,24^ have been constructed from their respective pan-genomes to investigate strain-specific adaptations and common functionalities. However, pan-genome models often require extensive manual curation and may encounter gap-filling and model consistency issues.

A second approach that is more streamlined involves reconstructing using existing curated species-specific metabolic models from resources such as AGORA^25,26^ and BiGG^27^. This approach addresses curation challenges and reduces the manual burden^21^, making it easier to scale up for the reconstruction of multiple genera. Prior studies have demonstrated the utility of the pan models created using the createPanModels routine from the Microbiome Modelling Toolbox (MMT)^28^, in studying the alteration in the metabolism of the human microbiome under different disease conditions^29–32^.

While there are no frameworks to consider the uncertainty in microbiome composition stemming from the inherent nature of 16S datasets, there are existing tools (‘createPanModels’) that can pool or concatenate multiple GEMs to create “pan” models. However, these models can not leverage the metagenomics-informed genus-specific data and often restrict their taxonomic input either to a genus or species level. Given that most amplicon sequencing data provides a combination of species and genus information, disregarding either of these components may impede accurate predictions of metabolic capabilities. To date, no existing tools have effectively addressed these ingrained complexities in microbiome composition data from 16S sequencing.

The current study proposes a new framework to generate a repository of flexible pan-genus metabolic models by harnessing the available curated GSMMs. The reconstructed PGMMs are validated for their applicability in modelling microbial communities with different levels of taxonomic information and exploring the metabolic intricacies within microbial constituents at the genus level. Our approach provides a way to surmount the inherent limited resolution of 16S rRNA sequencing and the limitations of current PGMMs in encapsulating the metabolic traits.

## 2 Methods

To mitigate the ambiguity stemming from 16S sequencing in metabolic modelling and bridge the gap in creating a species-aware pan-genus metabolic model (PGMM), the “Panera” method is proposed. The algorithm generates a pan-genus metabolic model from the existing strain/species-specific genome-scale metabolic models (GSMM) in the AGORA database. AGORA (Assembly of Gut Organisms through Reconstruction and Analysis) is a database of semi-automatically curated genome-scale metabolic reconstructions of human gut microbes. The AGORA database includes 818 metabolic reconstructions representing 1470 KEGG orthology identifiers (KO IDs), 227 genera and 14 different phyla. ModelSEED and KBase-based draft reconstructions of microorganisms from the annotated reference genome are gap-filled to ensure the reaction’s directionalities, mass, and balance charge. The gap-filled draft reconstructions are further refined with publications and comparative genomic analyses.

### 2.1 Formulation

Reconstruction of PGMM from species-specific GSMM of a selected genus was generated from the ‘createPanGenusModel.m’ in the Panera algorithm. The reconstruction pipeline employs three steps to produce a flexible PGMM: (i) Building a consensus model from the reactions in all the species of a genus, (ii) Formulating biomass to represent all the species in a genus model, and (iii) Adding fields to accommodate the variation in species composition. The steps included in the PGMM reconstruction are illustrated in Figure 1 and detailed in this section.

**Figure 1:**
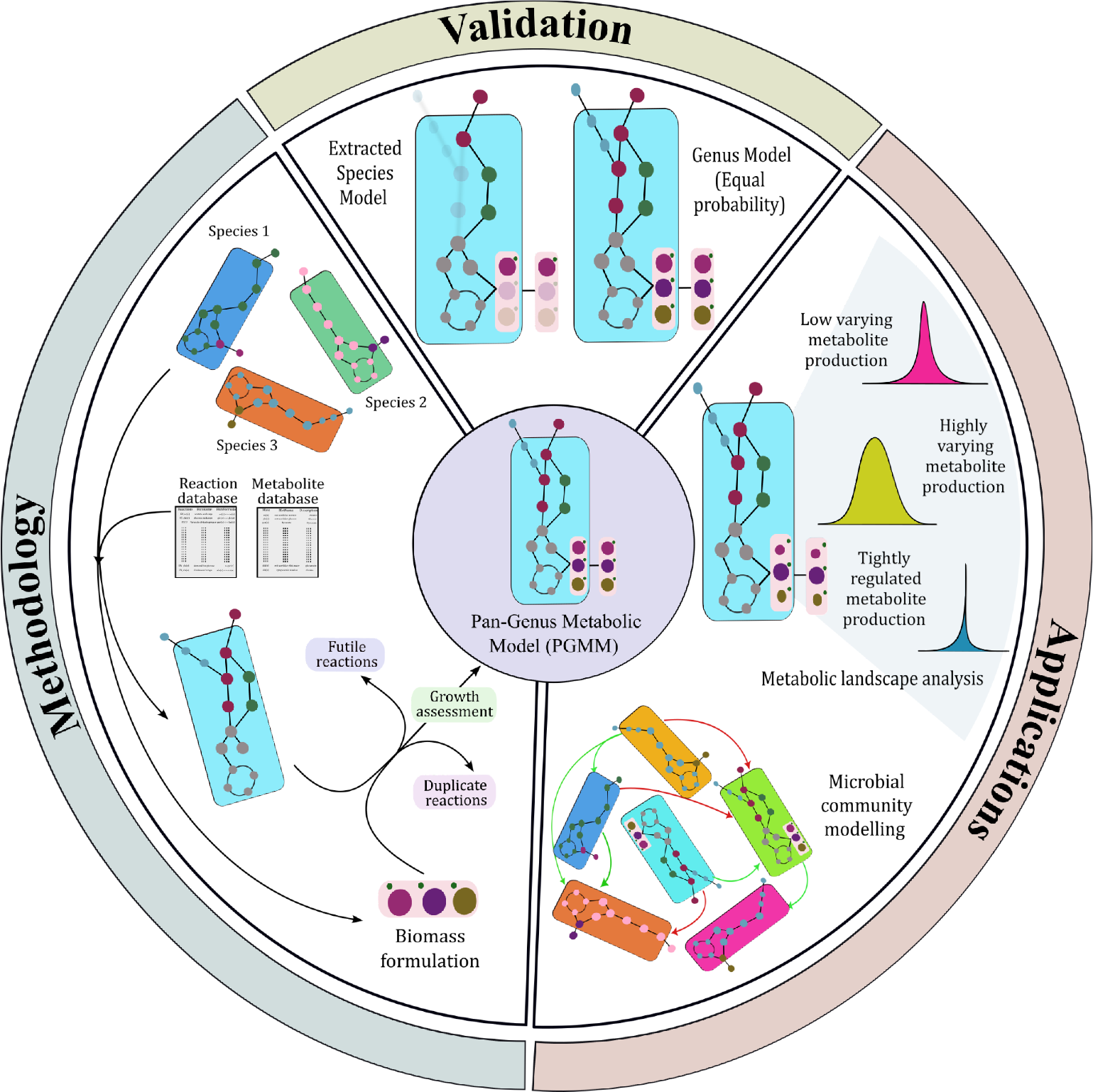
Methodology workflow of the Pan-Genus Metabolic Model (PGMM) reconstruction through ‘*Panera*’ methodology illustrated along with validation strategies and potential application of PGMM

#### 2.1.1 Building a consensus model from all the species GSMM of a genus

1. A database of all metabolites and reactions in Virtual Metabolic Human (VMH) models is retrieved from the Demeter pipeline^33^. A separate database for the biomass reactions and metabolites of the species models is generated for the reconstruction (Supplementary Table S1: Information of the species biomass reactions used in the model reconstruction).
2. Reactions from the GSMM models concerning a specific genus are extracted, and unique reactions (set of all the reactions) are selected to build a model.
3. Unique reactions, except species biomass reactions, are integrated into a model using rBioNet. The fields such as rxnNames (reaction names), grRules (gene reaction association), compNames (Compartment where the reaction takes place - cytosol or Extracellular) and subsystems are added using related information for the reactions from a reaction and metabolite database.

#### 2.1.2 Formulating biomass to represent the species in a genus model

4. The biomass reaction for the pan-genera model is formulated as the linear combination of biomass reactions of individual species in the genus:

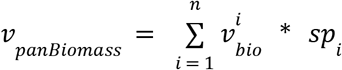

where *v*_*panBiomass*_ is the biomass flux of the pan-genera model (Objective function), *n* is the number of species in the genus, 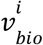 is the biomass flux of the *i*^*th*^ species and *sp*_*i*_ is the coefficient for *i*^*th*^ species, which implies the relative abundance or proportion of the microbial species in a community. The *sp*_*i*_ values can be adjusted to study the influence of a particular species in a genus by altering the species with different compositions. The reactions and metabolites associated with the ‘panBiomass’ and species biomass reactions are incorporated using biomass reaction and metabolite database. The default values of coefficients of species biomass (*sp*_*i*_) will be set to 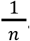. The default setting establishes an equal contribution from each species, and the coefficients can be adjusted to explore the distinct impact of a species.
5. Duplicate reactions or metabolites and reactions/metabolites involved in futile cycles are removed from the PGMM if the removal does not impact the growth of the model.
6. The refined pan-genus model is examined for growth by optimising the model with biomass as an objective while constraining to a particular media condition.

#### 2.1.3 Adding fields to accommodate the species composition variation

7. After PGMM refinement, a “reaction-species matrix”—a binary matrix representing whether the reaction is present (1) or absent (0) for an individual species - is combined as a field (‘rxnPresenceMat’) with the model.
8. An ‘spList’ field is incorporated into the model. Both ‘rxn-species matrix’ and ‘spList’ along with normalised ‘species probability vector’ will help filter the reactions to include in PGMM. The species probability vector is a vector of length nSp, representing the abundance of each species in the genus, where nSp represents the number of species in a genus).

### 2.2 Analysis of the reconstructed pan-genera metabolic model in analysing species metabolic abilities

The reconstructed PGMM, which represents the universe of reactions and metabolites present in all of the species within a genus, was used to perform *in silico* simulations. The PGMM was simulated to mimic the growth patterns of individual species and the collective growth of all species within PGMM to assess the fidelity of a reconstructed model. All the community model simulations were carried out using MATLAB R2022b. The figures were generated using the *ggplot2* package in R (version 4.0.1).

#### 2.2.1 Simulation of metabolic capabilities of an individual species

Initial simulations were carried out to explore the effect of including reactions from other species in PGMM while studying the metabolism of an individual species. We derived individual species models from PGMM using a species probability vector. To adapt the PGMM to specific species compositions, we adjusted the model based on the species probability/abundance. For instance, in the case of a PGMM representing a genus with five species, we simulated with a species probability vector indicating the presence of a single species at a time (Simulation 1: [1,0,0,0,0]; Simulation 2: [0,1,0,0,0] and so on).

Reactions with a reaction presence probability (product of species probability and reaction-species matrix) of more than zero were retained, while zero probability reactions were constrained by their lower and upper bounds. We adjusted the coefficients of the panBiomass reaction to represent a species model, and the model was subjected to Flux variability analysis^34^ (FVA). The maximum flux of FVA was used as an indicator for the metabolic production capability of the model. Additionally, we analysed the growth and maximum metabolite production potential of the species-specific GSMM. To evaluate the ability of PGMM to preserve the functionalities of a single species, we compared the metabolite production abilities between species-specific GSMM and modified PGMM under a given media condition. Jaccard distance between the maximum FVA values from GSMM and PGMM was evaluated to represent the qualitative variation by capturing the differences in the production of metabolites in the model, i.e. distinction in the metabolites with non-zero flux values. Whereas the Euclidean distance between the maximum FVA of PGMM and GSMM was calculated to explain the quantitative variation, i.e. the magnitude of variation in the production flux value of the metabolites in the model. This distance metric provides insight into how much the amount of produced metabolites differs between the models. To ensure comparability across different models, we normalised the Euclidean distance by dividing it by the maximum value observed among all the models.

#### 2.2.2 Working of Pan-genera metabolic model

The top-down approach of reconstructing PGMM aims to capture the genus-wide functionalities using species-level metabolic information. We assigned equal species probability as coefficients for biomass reactions in PGMM. For instance, in a genus with ten species, the coefficients for all the species biomass reactants in the panBiomass reaction were set to 0.1, representing equal contribution. We generated a reference multi-species community with an equal abundance of species within a genus using GSMM and Microbiome modelling toolbox (MMT). We performed flux variability of the microbial communities derived from PGMM and GSMM and compared the differences in the presence and magnitude of the metabolite production to gauge the ability of PGMM to represent the conserved and unique metabolic traits of a genus.

### 2.3 Application of PGMM in interpreting the metabolic landscape of the genus

The metabolic functional terrain of a genus could illustrate and cast light on the metabolic diversity trajectories^35^. We simulated the PGMM with varying species composition of a genus to explore their metabolic landscape. The varying species combination was implied on the model by applying a species probability vector, which was generated by normalising the sum of randomly generated values for each species within the genus to 1. By constraining the reaction bounds and species biomass coefficients in panBiomass, the model was tailored to the provided species composition.

Tailored PGMMs were subjected to FVA under different dietary conditions (Average European (EU) diet and Mediterranean diet (The constraints for the diet conditions were retrieved from VMH)) and utilised maximum flux from FVA to evaluate the metrics to define the flux bandwidth of the metabolites. Two different metrics, average maximum flux, which represents the mean of maximum flux of the metabolite production/consumption across different species composition and flux range, which explains the difference between the highest maximum flux to the lowest maximum flux observed for a metabolite across varying compositions were used to categorise the reactions into

i. No production - if both the averaged maximum flux and flux range are zero;
ii. low varying reactions - if the averaged flux is non-zero and the flux range falls between 5% to 50%;
iii. highly varying - if the flux range is greater than 50% and
iv. tightly regulated - if the flux range is within 5%.

### 2.4 Employment of PGMM in the analysis of microbial community metabolic modelling

To unravel the interplay between either the microbes in a community and the environment/host or the interaction within microbial species in an ecosystem, it is crucial to investigate their metabolic exchanges. Metabolic microbial communities constructed from metagenomics are valuable tools for probing the hidden complex microbial associations and interactions^36^. In the current study, we studied the utility of PGMM in a microbial community using synthetic and publicly available metagenome datasets. We aimed to understand how PGMMs compare to GSMMs in capturing community interactions in synthetic composition and real microbial metagenomic analysis.

#### 2.4.1 Application of PGMM in analysing metabolic capabilities of synthetic microbiota

##### 2.4.1.1 Synthetic abundance data generation

In this study, we generated synthetic abundance data for different community sizes, *k* (10 and 50), by randomly selecting ‘*k*’ strains from the pool of 818 AGORA metabolic reconstructions and assigning a random value to each strain. We performed data normalisation, ensuring that the total abundances summed to 1. The normalised data was then grouped at the genus level to construct a genus abundance matrix. Additionally, we explored *hybrid* models that use taxa information resolved at both species and genus levels. Specifically, we conducted simulations using abundance data where 55% of the taxa were resolved to the species level, while the remaining 45% were resolved only to the genus level.

##### 2.4.1.2 Microbiota models from synthetic abundance data

The generated abundance data were utilised to construct the personalised microbiota models. MMT creates a template community model comprising all the species and/or genera in the dataset. The personalised model was created by adjusting species or genus biomass coefficients in the community biomass equation. We built four different community model types using (i) GSMM, (ii) PGMM derived from the present work, (iii) Pan model created using createPanModel of the MMT (PanModel), and (iv) hybrid models, where both GSMM and PGMM were incorporated. We examined the functionalities of the communities against the GSMM-derived communities.

All the community models were constrained to the average European Diet, as reported in VMH^37^. The secretion and uptake flux of the exchange metabolites of diet-constrained community metabolic models were subjected to FVA. We conducted the computational analysis with a high-level, multi-process and high-performance method, ‘distributed FBA’ in Julia-1.6.8^38^, to accommodate the larger number of microbial members in a community. We estimated differences in metabolites with non-zero flux and variation in the magnitude of flux values of the metabolites between the community models using Jaccard and Euclidean distances, respectively. This assessment could shed light on the capacity of PGMM to encapsulate the species-model metabolic inference in the microbial community and to ascertain the advantages of the metagenomics-informed pan-genus model (PanGEM) over the lumped pan-genus model (MMT).

#### 2.4.2 Comparison of metabolic prediction of GSMM and PGMM with metabolomics data

The metagenomics data obtained from the gut microbiome of colorectal cancer (CRC) patients^39^ was employed to investigate the potential of PGMM in characterising the personalised metabolic microbial community. The study^39^ presented metabolomic and metagenomic information from the gut microbiome samples collected from healthy individuals and CRC patients. Among the 406 subjects sampled, we selected 15 samples to conduct a comparative analysis of microbial community functionality using various community modelling approaches in conjunction with metabolomics data. We preprocessed the normalised abundance values of microbial species in the selected samples by removing rare taxa, defined as taxa with an abundance lower than 1*10^−3^. Additionally, we converted a portion of the species information (approximately 45%) to the genus level, enriching the abundance table with both species and genus-level information, which mimics the taxonomic output of the amplicon sequencing data.

We constructed the personalised community models for each sample with different source models (GSMM, PGMM from our algorithm, PanModel from MMT and hybrid approach) using the processed abundance table. Subsequently, we comprehensively analysed these community models using ‘distributed FBA’ in Julia to assess their metabolite production potential. To gauge the accuracy of our predictions, we compared the production of secreted metabolites from the simulated microbial communities with the metabolites identified in the actual metabolomic data. For this comparison, we employed Jaccard and Euclidean distance metrics to evaluate the accuracy of our predictions and identify potential errors in predicting metabolic capabilities.

## 3 Results

Pan-genus metabolic Models (PGMM) reconstructed from species GSMM using the *‘Panera’* method aids in comprehending the functionalities of a microbial community when dealing with uncertain taxonomic information. The reconstructed PGMM comprises all the unique reactions from the species existing in the genus, as well as the metabolites and reaction-associated information from the metabolite and reaction databases (Supplementary Table S2: Metabolic information of the built PGMMs) along with the flexible biomass to incorporate prior species information. We validated the reconstructed PGMM by confirming their growth for a formulated biomass as an objective. The biomass formulation incorporates species biomass with their abundances, effectively reflecting the presence and the proportion of a microbial species in a sample. Our analysis of PGMM and hybrid models in microbial community models revealed that the hybrid models exhibited a noteworthy improvement of 10% in their ability to predict metabolic capabilities. This improvement underscores the efficacy of hybrid models in assessing microbial community models, particularly when compared to the lumped PanModel generated by MMT.

### 3.1 PGMM can be a representative of both genus and species

The ability of the PGMM to retain the functionalities of an individual species model was examined with a randomly selected microbial species from the AGORA database (Table 1: Information about the GSMMs used for the comparison). To evaluate the species representation ability of the PGMM from Panera, we compared their production/consumption potential of exchange metabolites with GSMM through FVA (Figure 2A). We considered a flux difference as significant if the ratio of the maximum flux of exchange metabolites in the GSMM to that in the species models from PGMM exceeded 10%. In qualitative analysis, which examines the number of metabolites produced within a model, we observed a minimal variation of only 0.01% to 2% between the models. In contrast, quantitative differences, which pertain to variations in metabolite production/consumption rates, showed a moderate variation of approximately 7-10% in the fluxes of exchange metabolites.

**Figure 2.**
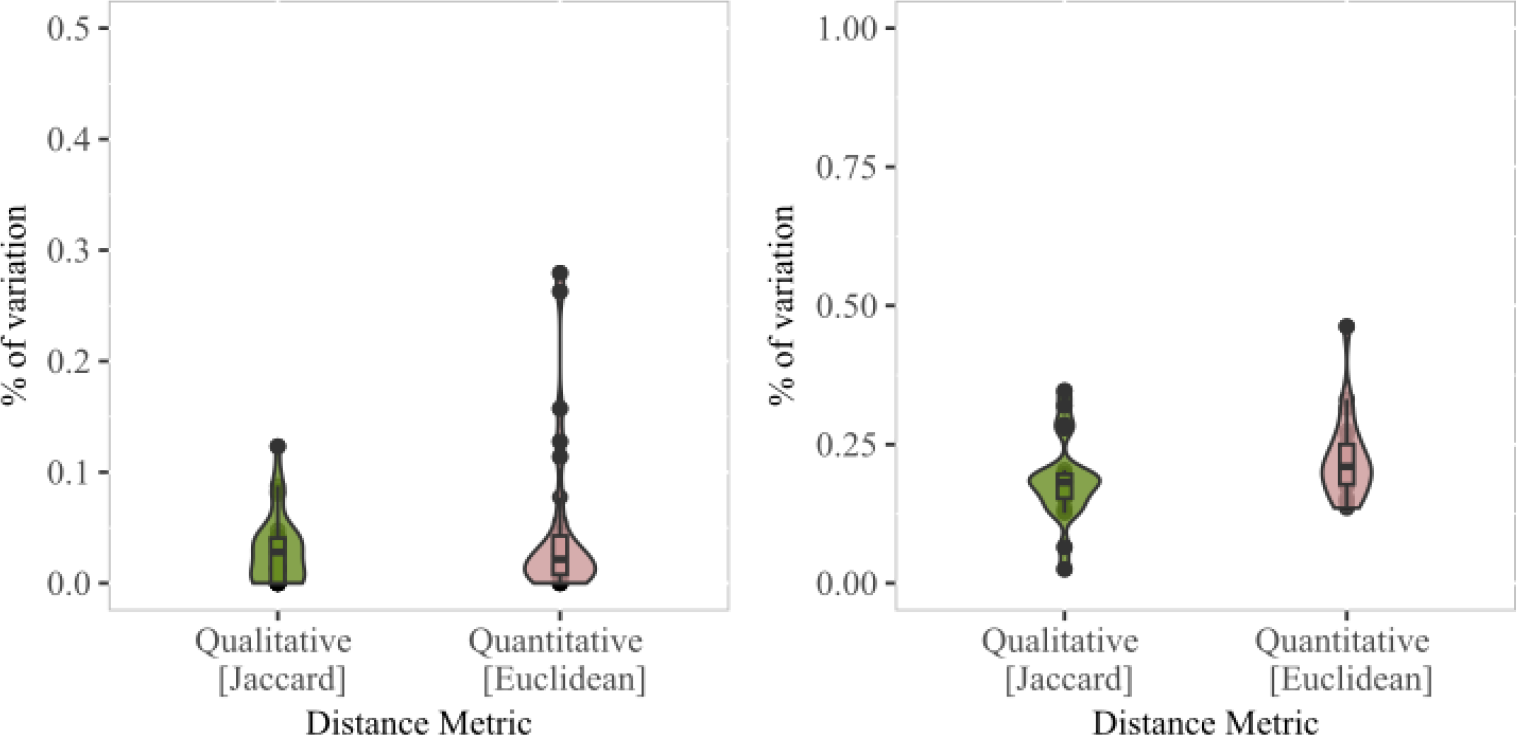
Qualitative and quantitative variation in the metabolic potential abilities for (A) extracted PGMM-Panera models in comparison to GSMM and (B) equal-species abundance implied PGMM in comparison to GSMM-based community models

The assessment of the potential of PGMM to represent the genus’s metabolic capabilities was carried out and validated by comparing it against the genus community models built using GSMMs. We applied a species probability vector in which the proportions are uniform and summed to 1 to revamp the PGMM to illustrate the equal presence of all the species within a genus. The genus community model was then reconstructed by integrating the GSMMs at equal abundances through compartmentalisation. In contrast to individual models, the genus representation by PGMM and GSMM-based communities showed substantial variability in both qualitative and quantitative analyses (Figure 2B). The analysis of maximum exchange metabolite production flux from the GSMM-based community model and tailored PGMM revealed an average difference of 22% in the production flux of exchange metabolites, along with a 17% variation in predicting the number of metabolites produced from the models.

### 3.2 Metabolic landscape analysis reveals a fascinating active metabolic similarity between genera

The flexible nature of the Panera-derived PGMM was leveraged to investigate the metabolic production bandwidth in a genus, which could help us understand the entire capability of a genus. We categorised the observed metabolic bandwidth into four different groups: high varying, low varying, tight regulation and no production. Tight regulation of specific metabolites in a genus, irrespective of the species composition, suggests that flux maintenance of the metabolite is conserved across all the species and crucial for the survival of the genus under the provided condition. Metabolic production with a larger flux range indicates the varying impact of individual species on the metabolite production potential.

Interestingly, the mapping of metabolic flux bandwidth across all the genera revealed that opportunistic pathogenic genera exhibit greater metabolic similarities among themselves than with the commensal genera (Supplementary Figure S2: Heatmap of the metabolic bandwidth for exchange metabolites across genera). This observation is supported by the distance tree built based on the similarity measure of metabolites (Supplementary Figure S1: Distance tree based on metabolites similarity of a genus) and reactions (Supplementary Figure S1: Distance tree based on reaction similarity of a genus), which rely on the presence or absence of the entity. These trees demonstrate a similar metabolic potential among the opportunistic pathogenic genera such as cluster-I (*Staphylococcus, Streptococcus, Shigella* and *Serratia*), cluster-II (*Haemophilus, Helicobacter, Klebsiella* and *Gemella*) and commensal genera (*Blautia, Bifidobacterium, Bacillus* and *Bacteroides*). Moreover, the flux bandwidth analysis provides insights into the characteristics of the genera. For instance, a distinct clustering of *Bacteroides* and *Prevotella* observed on the reaction and metabolite similarity-based trees is challenged by the flux bandwidth-based tree (Supplementary Figure S1: Distance tree based on the type of flux bandwidth observed under EU diet and Mediterranean diet), which proposes a shared metabolic regulation between the genera. A similar pattern was also observed for *Shigella, Enterobacter, Klebsiella* and *Haemophilus*. Remarkably, more than 70% of the clustering patterns using flux bandwidth types were found to be consistent across different dietary conditions (average European and Mediterranean diets).

In our analysis, we observed that the metabolic flux bandwidth of a specific metabolite exhibited variation among different genera, and this bandwidth seemed to be indirectly related to metabolite regulation. For example, we found that the flux range of L-Cysteine observed in *Bacteroides* and *Prevotella* was notably wider than the *Streptococcus* (Figure 4). However, regardless of whether the variability was high or low, we could discern the control mechanisms these different genera exerted over cysteine production through the bimodal distribution.

**Figure 4:**
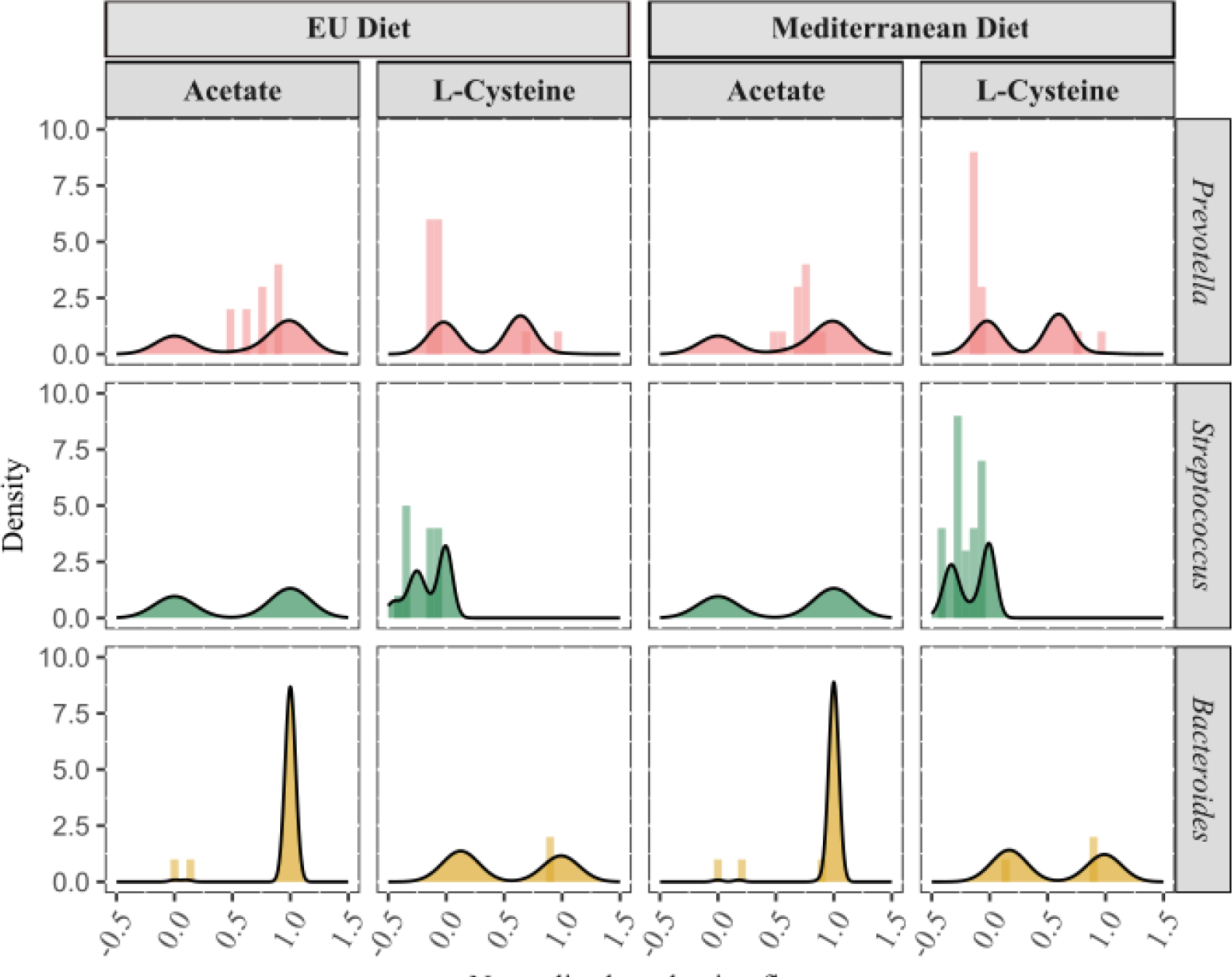
Metabolic flux bandwidth of acetate and L-cysteine production in a genus under European (EU) diet (left) and Mediterranean diet (right)

**Figure 4:**
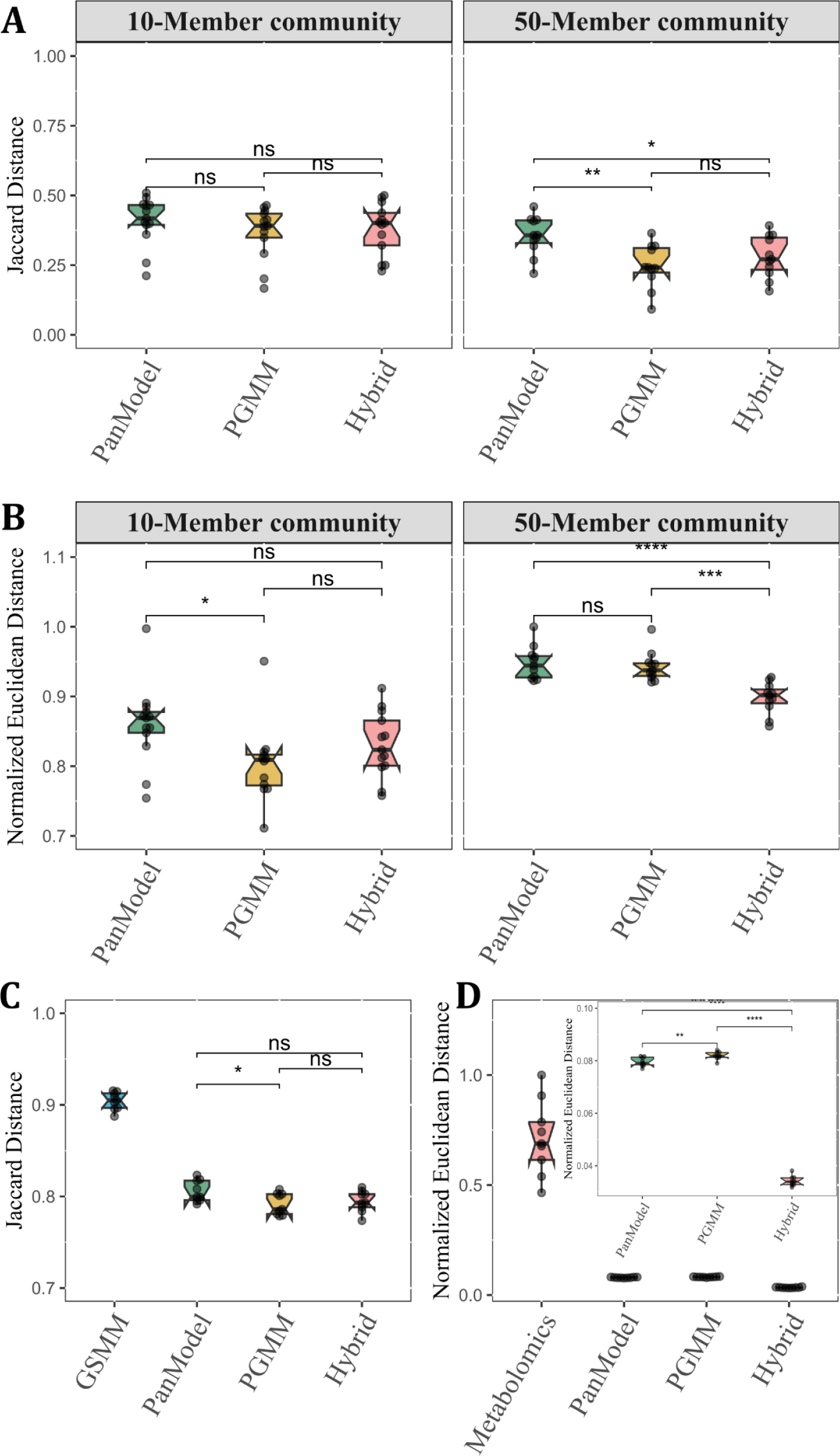
A comparative analysis of qualitative (Jaccard distance) and quantitative (Normalized Euclidean) differences is presented for two datasets: synthetic dataset (A and B) and CRC metagenomics dataset (C and D). The analysis encompasses various community types, including PGMM (Panera-derived Pan-Genus Metabolic Model), PanModel (Pan-Genus Metabolic Model constructed with ‘CreatePanModels’ in the CobraToolBox suite), and Hybrid communities (combining GSMM and PGMM). The values are calculated in relation to GSMM-based community models. (A) Highlights differences in metabolic diversity and (B) metabolite production when compared to GSMM-based communities, for both 10 and 50 microbial species communities in synthetic datasets. (C) Demonstrates the variance in metabolic production diversity within a community when compared to metabolomics study in the CRC metagenomics dataset. (D) Displays the differences in metabolic production flux within a community concerning GSMM-based communities in the CRC metagenomics dataset.

Furthermore, the fluctuating flux range distribution for acetate and L-cysteine across these three genera highlighted the control implementation depicted by the species composition. Upon deeper investigation into the metabolic flux bandwidth of each exchange metabolite for these genera (Supplementary Figure S2), we discovered that metabolites linked to inorganic ion metabolism, like Zinc, Copper, and Magnesium, exhibited more stringent regulation across different genera. This may be attributed to the limited need for micronutrients in the survival of these organisms.

We observed that the cofactor group, which includes quinone, glutathione, folate and riboflavin, showed a robust regulation in most of the genera, except for the genus cluster containing *Shigella, Escherichia, Enterococcus, Klebsiella* and *Haemophilus* under both dietary conditions. On the other hand, this specific genus cluster displayed robust regulation for other cofactors like reduced glutathione. This suggests that different genera possess unique requirements for cofactors, which is crucial for their functional activities. In contrast, amino acid production fluxes, another class of metabolites, exhibited greater variability than the other metabolites group. Notably, commensals displayed a weak regulation over amino acid production compared to other genera. Most of the observed results remained consistent under both dietary conditions used for the simulations. Consequently, we detected higher similarity in metabolite-level clustering between EU and Mediterranean diet conditions.

### 3.3 PGMM captures better metabolic information than the lumped genus model in a community modelling

Synthetic microbial abundance data for different community sizes (*n* = 10 and 50) was used to evaluate the applicability of PGMM in characterising the microbial communities through metabolic modelling. We reconstructed four different community types and compared their metabolic output with the widely used GSMM-based community outcomes. The qualitative variation analysis revealed that Panera-derived PGMM models encapsulated more metabolic information than Pan models when employed in communities, and this difference became highly significant with an increase in community size (Figure 4A).

Furthermore, the quantitative variation analysis explained that PGMM-based communities predicted the measure of metabolic capabilities more accurately than the Pan model-based communities compared to the GSMM-based microbial communities (Figure 4B). However, the uptake fluxes, i.e., the consumption capability of a community, did not show any significant variation (Supplementary Figure S3). This analysis also suggested that even a smaller variation in the nutrient uptake in the model results in substantial changes in production fluxes, possibly due to the modification in the model structure of each community model. In conclusion, the findings indicated that PGMM communities have the ability to project GSMM-based communities and could serve as a valuable alternative to GSMM for community metabolic modelling with ambiguous taxonomic data.

### 3.4 Improved prediction of metabolic potential using hybrid models

The microbial communities generated with the CRC-metagenomics data were subjected to qualitative and quantitative variation between the models and compared to the available metabolomics. We assessed the qualitative variation by comparing the metabolites detected in the metabolomics data with those produced from the various community types (Figure 4C). Interestingly, a greater number of secreted metabolites was captured by the communities built from PGMM, Pan models, and hybrid models (GSMM along with PGMM), than by the GSMM-based microbial communities. Notably, the metabolic capacity of over 25 metabolites, including amino acids and secondary metabolites like cholate derivatives, was better represented than GSMM.

To quantitatively evaluate the alteration in the production flux of exchange metabolites, we reported the comparison with GSMM-based communities since the metabolomics data were presented in terms of metabolite concentration. Despite the observed significant differences between PGMM and Pan model communities, the magnitude of variation only fell within the range of 0.2 to 0.5% (Figure 4D). Moreover, hybrid community models (using both GSMM and PGMM to build a community) demonstrated a nearly 50% reduction in error rate in functionality prediction. Overall, the evaluation of metabolomic analysis outcomes against the predicted metabolic abilities of the various community models insinuates that hybrid models exhibit better prediction accuracy with minimal qualitative and quantitative disparities.

## 4 Discussion

Metagenomic sequencing technologies, particularly the cost-effective 16S rRNA sequencing, have significantly advanced our comprehension of microbial ecosystems, specifically regarding their compositional and functional dynamics. Nevertheless, the fundamental limitation of acquiring taxonomic assignments at finer taxonomic levels in amplicon sequencing poses a challenge in functionality prediction using constraint-based *in-silico* microbial community modelling. To address the issue, the Pan-Genus Metabolic Model (PGMM) is employed as a valuable alternative to Genome-Scale Metabolic Models (GSMMs) for constructing microbial communities and studying their functionalities and dynamics. However, reconstructing high-quality PGMMs from pan-genomes requires time-consuming manual curation and has led to the exploration of an alternative approach using curated GSMMs. Yet, PGMMs generated from existing tools, such as the Microbiome Modelling Toolbox (MMT), suffer from limitations in accommodating both the genus and species compositionality and representing the species within a genus, which restricts their utilisation in community modelling that incorporates both genus and species information from amplicon sequencing. Consequently, the full potential of PGMMs for microbial community analysis remains underutilised.

To address these challenges, we introduce our novel method, “*Panera*”, which presents a unique and adaptable framework for generating PGMMs. The primary aim of Panera is to reduce uncertainties in assessing the metabolic capabilities of a microbial community while using 16S amplicon sequencing data. In this framework, we construct a comprehensive model by integrating all unique reactions from individual species-specific GSMMs within a genus and their respective metabolic data. The ‘panBiomass’ equation, which represents a linear combination of species biomass equations, is then incorporated into the model to obtain a PGMM containing all reactions and pathways from the GSMMs. The components in PGMM are species-aware in contrast to the pan-genus models built from MMT. The Panera algorithm further enhances PGMMs by improving biomass formulation and introducing flexibility, allowing users to tailor the model to the specific input parameters. We rigorously tested Panera models to predict the metabolic capabilities of individual species and the collective abilities of genera. These models were evaluated for their applicability in exploring the metabolic landscape of genera and simulating *in-silico* microbial communities, particularly in comparison to conventional GSMMs.

Our analyses demonstrated that PGMMs effectively capture the metabolic activities of individual species GSMMs, as indicated by comparable qualitative and quantitative metabolic flux predictions between the two models (Figure 1). However, disparities arose when comparing PGMMs to GSMM-based microbial communities during simulations of equal species probability genus models. These discrepancies could be attributed to structural differences between the models. PGMMs exclusively comprise unique reactions within a genus in a single compartment, while GSMM-based communities adopt a compartmentalisation approach^40,41^ that combines all species models via an extracellular compartment. Despite both models utilising a similar objective formulation, which involves a linear combination of species proportion and species biomass, variations emerged when simulating a community biomass flux. A single unique biomass precursor reaction accounts for the multiple species biomass production in PGMM. In comparison, the GSMM-based community relies on precursor reactions within each species for their respective biomass production. Differences also surfaced in the prediction of certain biomass precursors, including higher fluxes for amino acids like cysteine and phenylalanine in PGMMs and lower fluxes for secondary metabolites such as cholate and phenol, alongside polysaccharide precursors (N-Acetyl D-glucosamine and glucosamine). These variations could be attributed to a potential trade-off between accuracy and abstraction stemming from information loss^42^.

We further investigated the potential of PGMMs in exploring the metabolic landscape of the genus. We simulated PGMMs with distinct species probability vectors, replicating varying species proportions in a genus. The analysis of reaction and metabolite similarity between PGMMs illustrated the clustering of genera with similar functions compared to the phylogenetic tree, aligning with previous findings indicating shared functionalities among phylogenetically diverse organisms^43–45^. Notable disparities between the reaction and metabolite similarity tree and metabolic flux bandwidth distance tree suggested that active metabolic fluxes may differ based on nutritional supplements and environmental factors compared to the potential functionalities observed within genera^35,46^. Additionally, the clustering of *Prevotella* and *Bacteroides* in the flux bandwidth-based tree could be supported by the shared core protein similarity^47,48^ between the two genera despite their associations with different diets^49^. Similarly, metabolic regulation clustering observed among opportunistic pathogens such as *Escherichia* and *Shigella* is consistent with their genetic similarity^50,51^. Moreover, significant variations in the amino acid production potential of dysbiotic communities^31,52^ present evidence for the enhanced regulation of amino acid production in opportunistic pathogens. Ultimately, the flexible Panera PGMM proved to be a valuable resource for investigating the capabilities of microbial genera and customising species composition within PGMMs provides a significant advantage for studying core functionalities and niche development.

However, a major limitation in constructing PGMMs from GSMMs lies in the quality of the PGMM, which is contingent upon the quality of source models. Additionally, these GSMMs should have a consistent standard annotation to ensure that the combined reactions function seamlessly as a proper metabolic model. This limitation constrains using the *‘Panera’* algorithm to models obtained from a single source with uniform annotation. Nevertheless, the current availability of 7302 curated strain-specific metabolic reconstructions, comprising 504 genera in AGORA2^26^, presents a substantial resource for PGMM reconstruction.

Finally, we assessed the principal utility of PGMM, which is to act as a valuable tool in constructing microbial communities with incomplete taxonomic information. The evaluation of PGMM in a microbial community used two distinct datasets: (i) synthetic microbial abundance data with different community sizes and (ii) metagenomics and metabolomics data collected from healthy individuals and CRC patients. Both analyses revealed that hybrid community models, which incorporate both GSMMs and PGMMs, offer predictions comparable to GSMM communities, surpassing the performance of PGMM or Pan model communities. As expected, PGMM communities outperformed Pan model communities across both datasets. The predictability of hybrid models was particularly efficient with larger community sizes, demonstrating nearly a 10% metabolic flux prediction reduction in 50-member community models. Notably, previous studies often simplified 16S rRNA taxonomic information to the genus level for metabolic analysis^31,32^ can use hybrid model communities as a promising alternative without compromising data richness. This strategy is especially pertinent since 16S rRNA sequencing provides a combination of species and genus information. PGMMs can be tailored to incorporate prior probabilities if the information is available for a better accurate representation of a genus in a given environment. For example, suppose the adult gut microbiota is known to comprise 50% *Bacteroides fragilis*, 30% *Bacteroides vulgatus* and 20% of the remaining species within *Bacteroides* genus. In that case, these prior probabilities can be applied to create a more precise model of the *Bacteroides* in the gut microbiota. Despite the variations in metabolic predictions, the adaptable PGMM and hybrid GSMM-PGMM communities demonstrate their significance in studying the metabolic abilities of microbial communities reconstructed from ambiguous amplicon sequencing data.

In summary, we have developed a novel framework, “Panera”, which can significantly reduce uncertainties in metabolic profiling of personalised microbial communities using relative abundance data obtained from 16S rRNA sequencing analysis. The flexible nature of the PGMM facilitates the analysis of metabolic profiles at varying species compositions within a genus, enabling the exploration of the metabolic landscape of genera. Furthermore, our study demonstrates that the hybrid community model, combining PGMMs and GSMMs, is a viable and efficient approach for capturing the capabilities of a microbial community, even when faced with uncertain taxonomic information.

## 5 Supplementary Information

Table S1: Information on the species biomass reactions used in the model reconstruction Table S2: Metabolic information of the built PGMMs

Figure S1: Distance trees based on similarity between reactions, metabolites and flux bandwidth across genera

Figure S2: Heatmap of metabolic flux bandwidth of each reaction in the model across multiple genera under average European and Mediterranean diet conditions

Figure S3: Boxplot of qualitative and quantitative variation in the uptake flux of the exchange metabolites in microbial communities

## 6 Acknowledgements

The authors thank Maziya Ibrahim for useful comments on the manuscript and Pratyay Sengupta for the help with the figures.

## 7 Code availability

The code for the PGMM construction using *‘Panera’* methodology is available at: https://github.com/RamanLab/Panera/

