## Supplementary Figures for "Panera: A novel framework for surmounting uncertainty in microbial community modelling using Pan-genera metabolic models"

### Supplementary Information

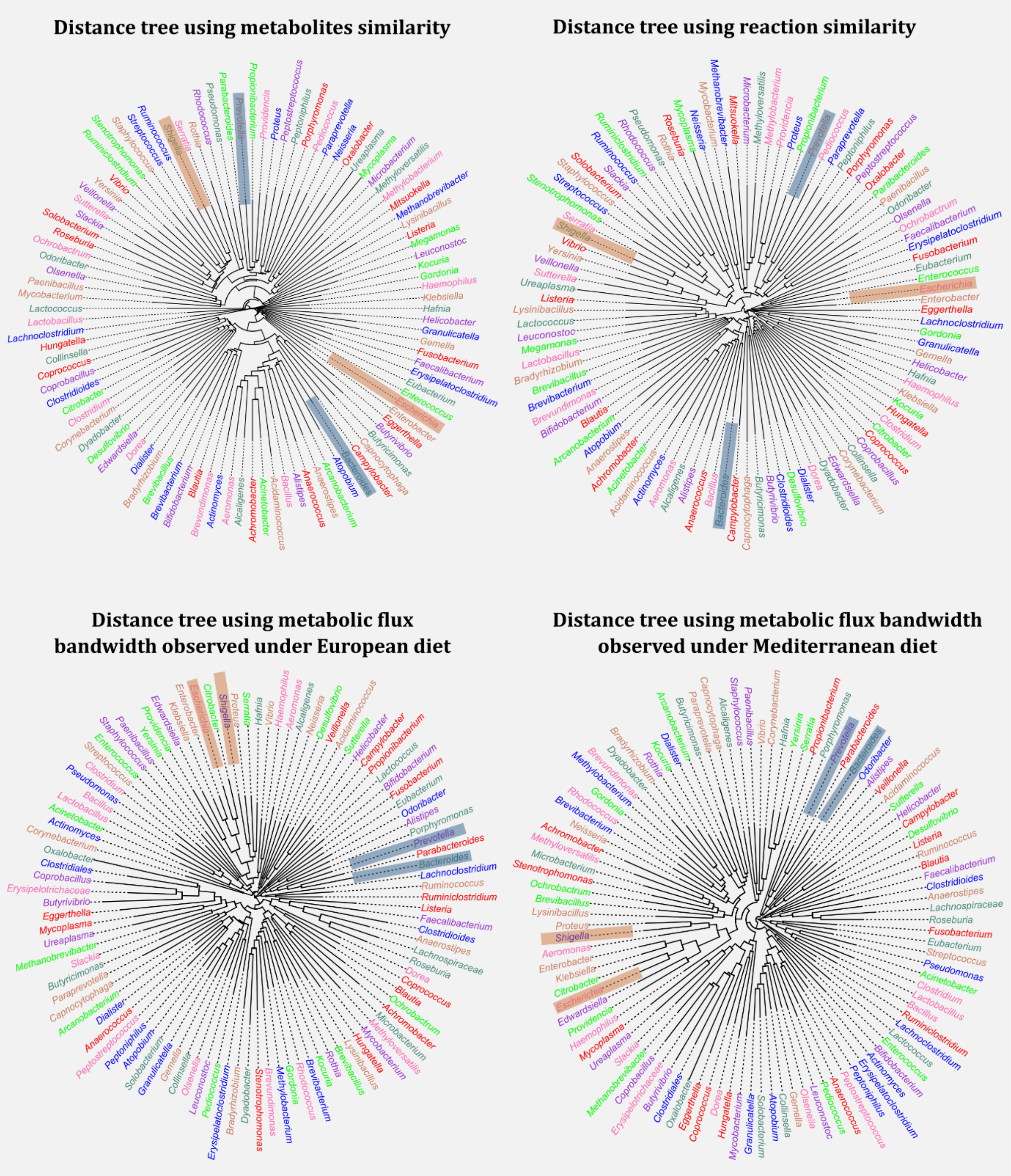


**Supplementary Figure S1:** Distance tree based on the Jaccard similarity depicting the relationships between genera regarding reaction, metabolite, and flux bandwidth types in various diets. The colour of the text within the tree indicates the phylum of each genus.

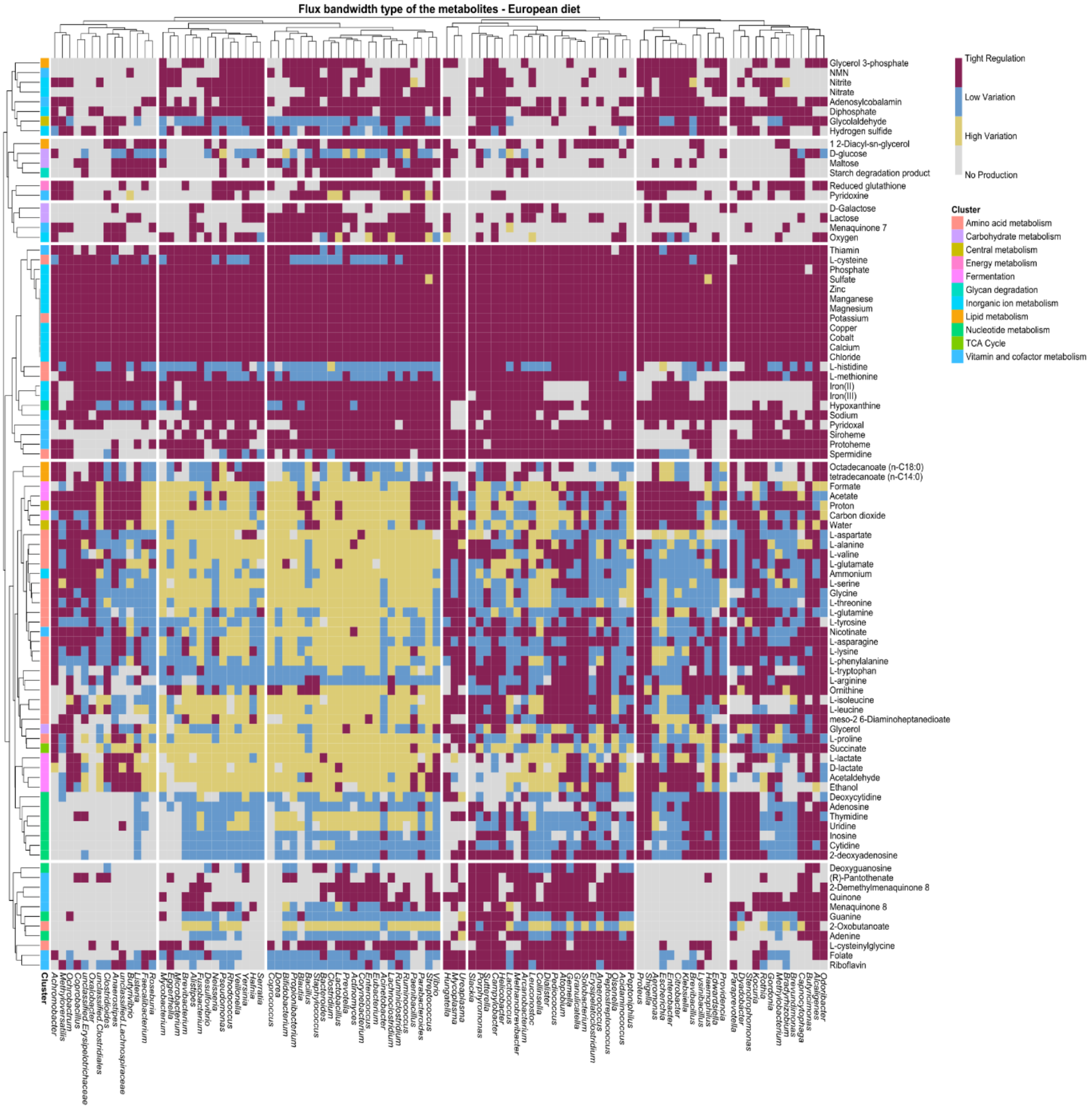


**Supplementary Figure S2a:** Heatmap illustrates the bandwidth patterns of exchange metabolite production across genera for a European diet. In this representation, rows correspond to exchange metabolites, and columns depict the individual genera. The figure excludes the metabolites which are not produced in most of the genera.

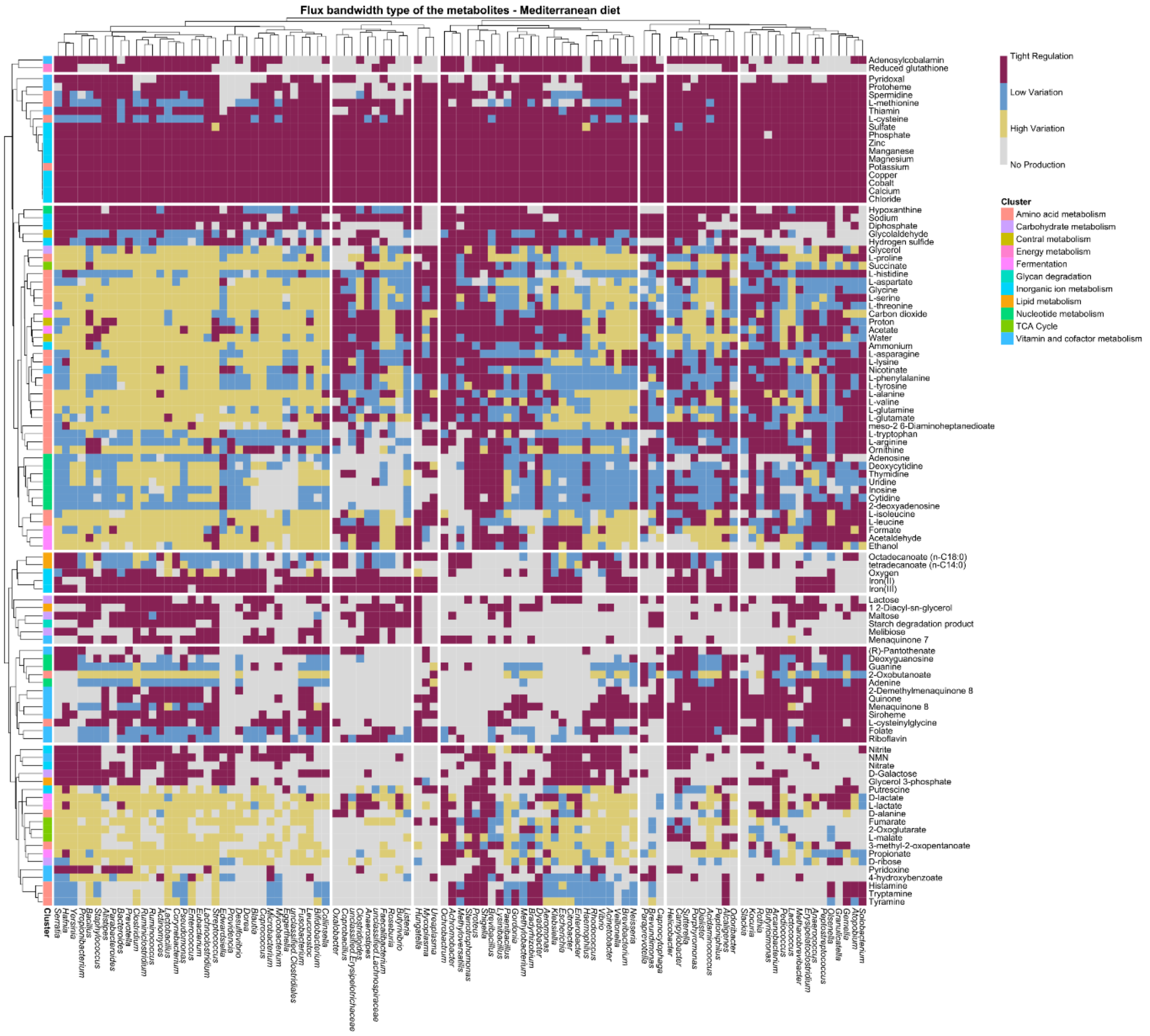


**Supplementary Figure S2a:** Heatmap illustrates the bandwidth patterns of exchange metabolite production across genera for a Mediterranean diet. In this representation, rows correspond to exchange metabolites, and columns depict the individual genera. The figure excludes the metabolites which are not produced in most of the genera.

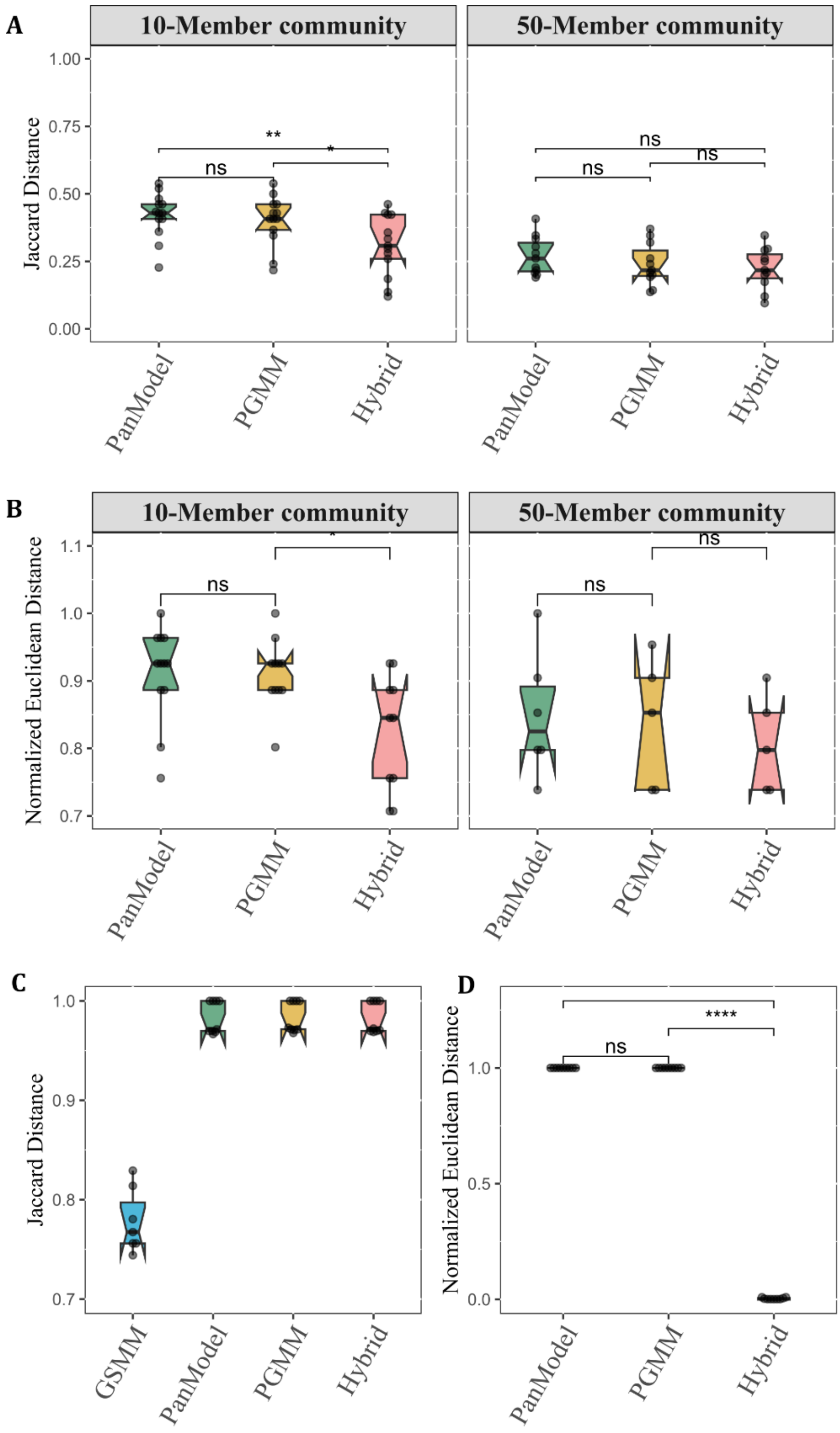


**Supplementary Figure S3:** A comparative analysis of qualitative (Jaccard distance) and quantitative (Normalized Euclidean) differences is presented for two datasets: synthetic dataset (A and B) and CRC metagenomics dataset (C and D). The analysis encompasses various community types, including PGMM (Panera-derived Pan-Genus Metabolic Model), PanModel (Pan-Genus Metabolic Model constructed with 'CreatePanModels' in the CobraToolBox suite), and Hybrid communities (combining GSMM and PGMM). The values are calculated in relation to GSMM-based community models.

(A) Highlights differences in metabolic diversity and (B) metabolite production when compared to GSMM-based communities, for both 10 and 50 microbial species communities in synthetic datasets.
(C) Demonstrates the variance in metabolic production diversity within a community when compared to metabolomics study in the CRC metagenomics dataset.
(D) Displays the differences in metabolic production flux within a community concerning GSMM-based communities in the CRC metagenomics dataset.
